# The effect of individual learning on collective foraging in honey bees in complex environments

**DOI:** 10.1101/817270

**Authors:** Natalie J. Lemanski, Chelsea N. Cook, Cahit Ozturk, Brian H. Smith, Noa Pinter-Wollman

**Affiliations:** Department of Ecology and Evolutionary Biology, University of California, Los Angeles, CA, 90095; School of Life Sciences, Arizona State University, Phoenix, AZ, 85287

**Author notes:** **Corresponding Author:** Natalie J. Lemanski; Department of Ecology and Evolutionary Biology, Life Sciences Building, University of California, Los Angeles, CA, 90095.

**Keywords:** Collective Behavior, Exploration-Exploitation Trade-Off, Foraging, Honey Bee, Latent Inhibition, Learning

## Abstract

The trade-off between exploiting known resources and exploring for new ones is a complex decision-making challenge, particularly when resource patches are variable in quality and heterogeneously distributed in the landscape. Social insect colonies navigate this challenge, in the absence of centralized control, by allocating different individuals to each of these tasks based on variation in individual behavior. To investigate how heritable differences in individual learning affect a colony’s collective ability to locate and choose among different quality food resources, we develop an agent based model and test its predictions using two genetic lines of honey bees, selected for differences in their learning behavior. Here we show that, paradoxically, colonies containing individuals that are better at learning to ignore unrewarding stimuli are worse at choosing the highest quality resource at the collective level. This work highlights the importance of individual variation within groups on the emergence of collective behavior.

## Introduction

Fitness strongly depends on an animal’s ability to find resources. However, animals often face trade-offs between exploiting known resources and exploring for new ones. How animals resolve this trade-off is influenced by the characteristics of the environment, such as how resources are distributed spatially in the landscape [1–4] and the variability of resource quality [5–7]. If most resource patches are similar in quality, organisms should persist in known patches, as long as those remain sufficiently profitable [7], because exploration is unlikely to yield something better. However, if resource patches are highly variable in quality, it could be beneficial to invest in exploring the environment to increase the chances of finding and exploiting the most rewarding resources.

Animals living in social groups, such as social insects, have the unique ability to engage in exploration and exploitation simultaneously by allocating each of these tasks to different individuals. In eusocial insects, the colony is the unit of selection because workers are sterile. How a colony distributes its foragers between exploration and exploitation can determine the collective foraging success of the entire colony and therefore its reproductive success [4,8,9].

Just like solitary animals, a colony’s investment in exploration and exploitation should depend on characteristics of the resource landscape. Previous theoretical work predicts that small resource patches, high patch density, and low search costs should increase a social group’s optimal investment in exploring [9]. Other work suggests that the benefits of exploitation through social recruitment, are highest when resources are difficult to find, patchily distributed, and variable in quality [10–12]. Colonies that invest more in exploitation (e.g., via recruiting) should therefore perform better when resources are clumped in the landscape, while colonies that invest more in exploration (e.g., via scouting) should perform better when resources of variable qualities are evenly distributed throughout the landscape. However, it is not known how the spatial distribution of resources influences a colony’s ability to choose the best resources, when those resources vary in quality.

Colonies of honey bees are composed of individuals that differ in whether they explore or exploit the environment. These differences in colony composition can have important consequences for collective foraging [8]. Among the foragers of a colony, a small fraction are scouts, who specialize on searching for new food resources, while the rest of the foragers exploit these resources [13,14]. The higher quality a discovered resource is, the longer a scout will spend recruiting other foragers to it [15–17]. Although it is not fully understood what causes certain individuals to act as scouts [18–22], the allocation of foragers to scouting or recruiting is influenced by variation among individuals within a colony in learning behavior, brain amine production, and gene expression [20,21,23].

A learning behavior that has been linked with honey bee foraging is latent inhibition, which is the tendency of an individual to ignore stimuli that have been repeatedly encountered without a reward [24]. Recent work has shown that scouts tend to be higher in latent inhibition than other foragers [23]. The tendency to ignore previously unrewarding ‘familiar’ stimuli may help scouts seek out new food resources when known patches begin to deplete, while lower latent inhibition may allow recruits to continue exploiting depleting patches until new ones are located [8,23]. Individual differences in expression of latent inhibition are heritable in honey bees and therefore there is natural variation in latent inhibition within a colony because of the genetic diversity that results from a queen mating with multiple drones [25]. Latent inhibition is exhibited by drones and queens as well as by workers, making it possible to artificially select lineages of honey bees that are higher or lower than average in latent inhibition [25]. Workers from these artificially selected lines display similar latent inhibition to their parents, regardless of adult social environment [26]. This gives us the ability to experimentally manipulate the composition of colonies to explore how individual learning affects colony level foraging behavior in different environmental situations.

Here we examine how latent inhibition (LI) affects the foraging behavior of honey bee colonies in differently structured landscapes. When resource patches are variable in quality, we expect the allocation of workers to exploration or exploitation to affect a colony’s ability to find the highest quality resource patch. Because high LI bees are more likely than low LI bees to act as explorers, we predict that colonies composed of high LI individuals would be better at finding the best quality resources. Because low LI individuals are more likely to act as exploiters, we predict that colonies composed of mostly low LI individuals would exploit effectively all known resources, regardless of their quality. Furthermore, optimal foraging theory suggests that colonies that invest more in exploration should be better at finding evenly dispersed resources, while colonies that rely more on recruitment should be better at exploiting clumped resources. We therefore predicted that colonies composed of mostly high LI individuals would perform best when resources are evenly distributed in the environment while colonies composed of mostly low LI individuals would perform best when resources are concentrated in a few large patches.

To answer these questions, we first developed an agent-based model to explore the effects of environmental features on the foraging behavior of honey bee colonies composed of a wide range of ratios of exploring and exploiting individuals (Figure 1A). We then tested our model predictions empirically by placing honey bee colonies that were genetically selected for either high or low LI in different environmental conditions that differed in the distribution of resources that varied in quality (Figure 1B).

**Figure 1.**
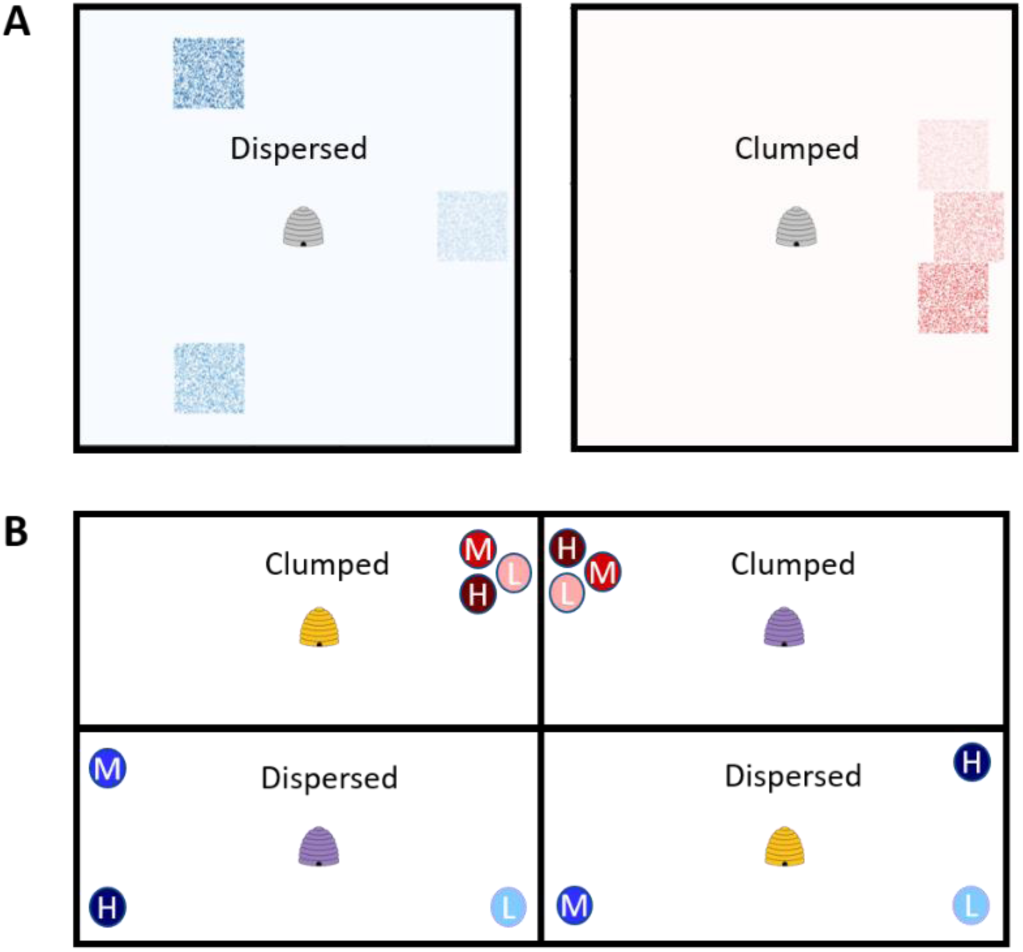
Spatial distribution of resources patches of different qualities in the (A) agent based model and (B) empirical experiment. (A) Simulated resource landscapes in the agent based model, with the symbol in the center indicating the hive. Squares are resource patches, with darker hues indicating higher quality. (B) Experimental setup of colonies in the empirical study. Each compartment (large rectangle) is a flight tent. Colored symbols in the middle of each tent depict a bee hive. Color indicates whether the colony is from a high (yellow) or low (purple) LI genetic line. Circles indicate feeders. Clumped treatment is in red and dispersed treatment in blue. After two days in this configuration, the experimental set up was flipped such that the top two flight tents received dispersed feeders in the open corners of the tent and the bottom two tents received clumped feeders in the open corner of the tent. Intensity of circle color and the letter inside it indicate feeder quality (H: high (2.5M), M: medium (1.5M), and L: low (0.75M)).

## Results

### Agent-Based Model

In our simulations, the proportion of scouts in a colony affected the colony’s ability to distinguish between resources of different qualities. Colonies with a lower proportion of scouts were better at choosing the highest quality food source than colonies with a high proportion of scouts (Figure 2). Regardless of resource distribution, colonies with a low proportion of scouts collected more food on average than colonies with a high proportion of scouts, but with higher variance (Figure 3). Furthermore, regardless of the proportion of scouts, simulated colonies were better at differentially exploiting the highest quality food source when resources were clumped than when resources were dispersed (Figure 2). Thus, contrary to our initial intuition, our model predicted that if high LI workers are more likely than low LI workers to behave as scouts, low LI colonies should be better than high LI colonies at finding and exploiting the highest quality feeder.

**Figure 2.**
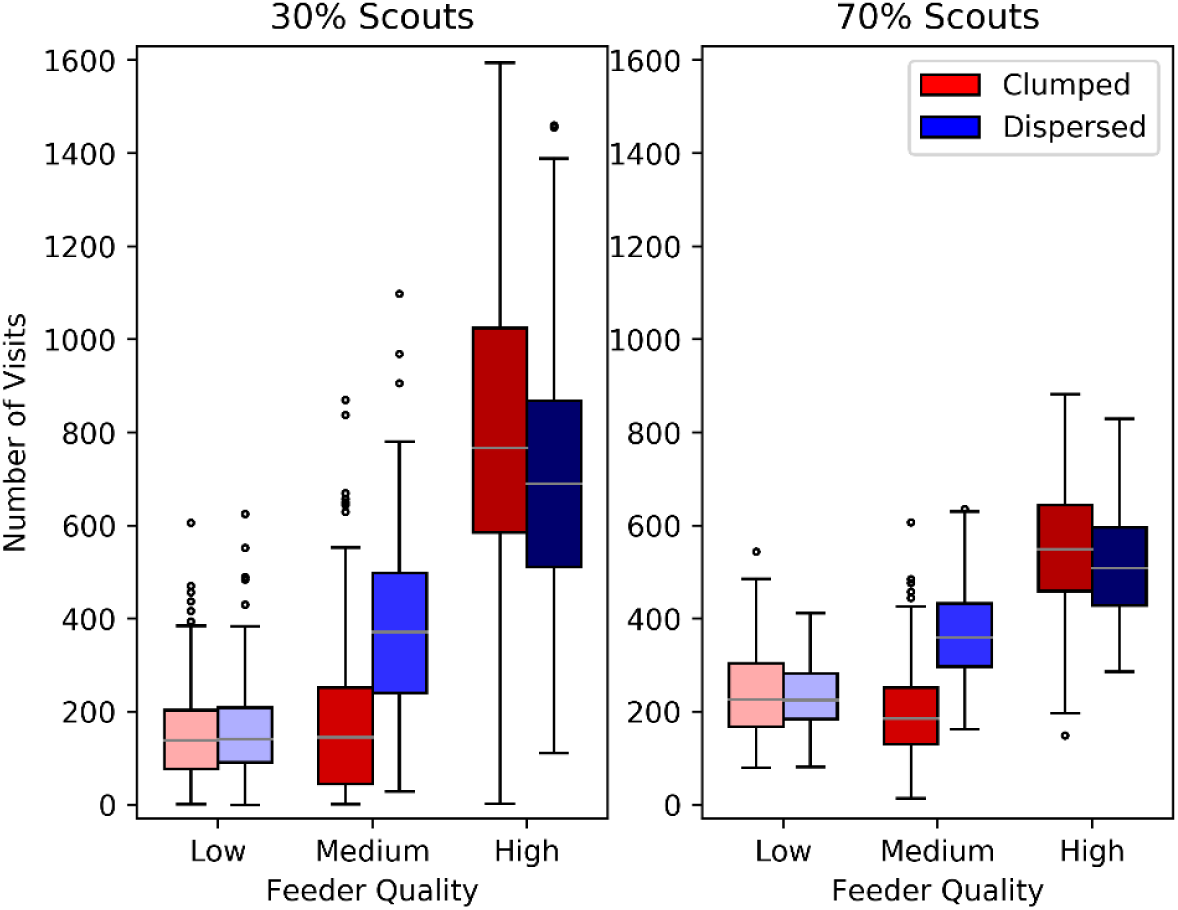
Simulated foraging behavior of colonies with low and high proportion of scouts. Colonies with (A) low proportion (30%) or (B) high proportion (70%) of scouts foraged on clumped (red) or dispersed (blue) resources that were of low (light hue), medium (medium hue), or high (dark hue) quality. Here and in all following boxplots, horizontal lines are the medians, boxes and whiskers indicate first and second quartiles, and dots are outliers.

**Figure 3.**
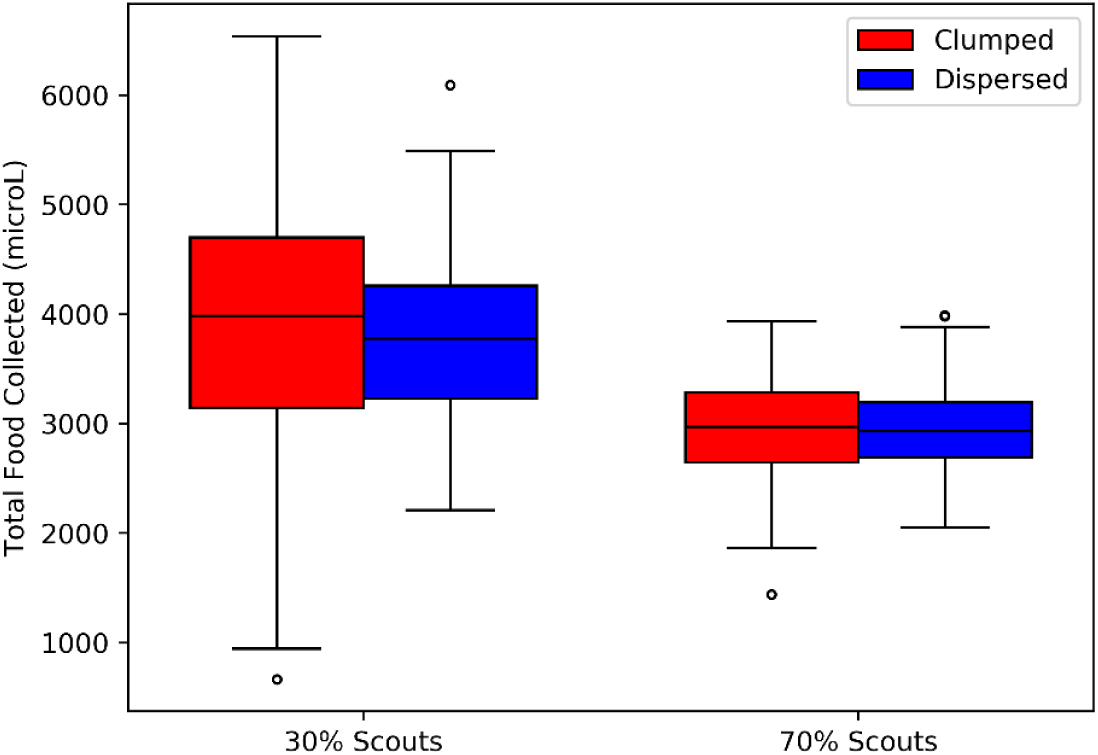
Total amount of simulated food collected. By colonies with low (30%) and high (70%) proportion of scouts when resources were clumped (red) or dispersed (blue).

Our model revealed that the spatial distribution of resources only weakly influenced the optimal investment in scouting. Contrary to our expectations, the optimal proportion of scouts was higher when resources were clumped (∼40%) than when they dispersed (∼30%) (Figure 4).

**Figure 4.**
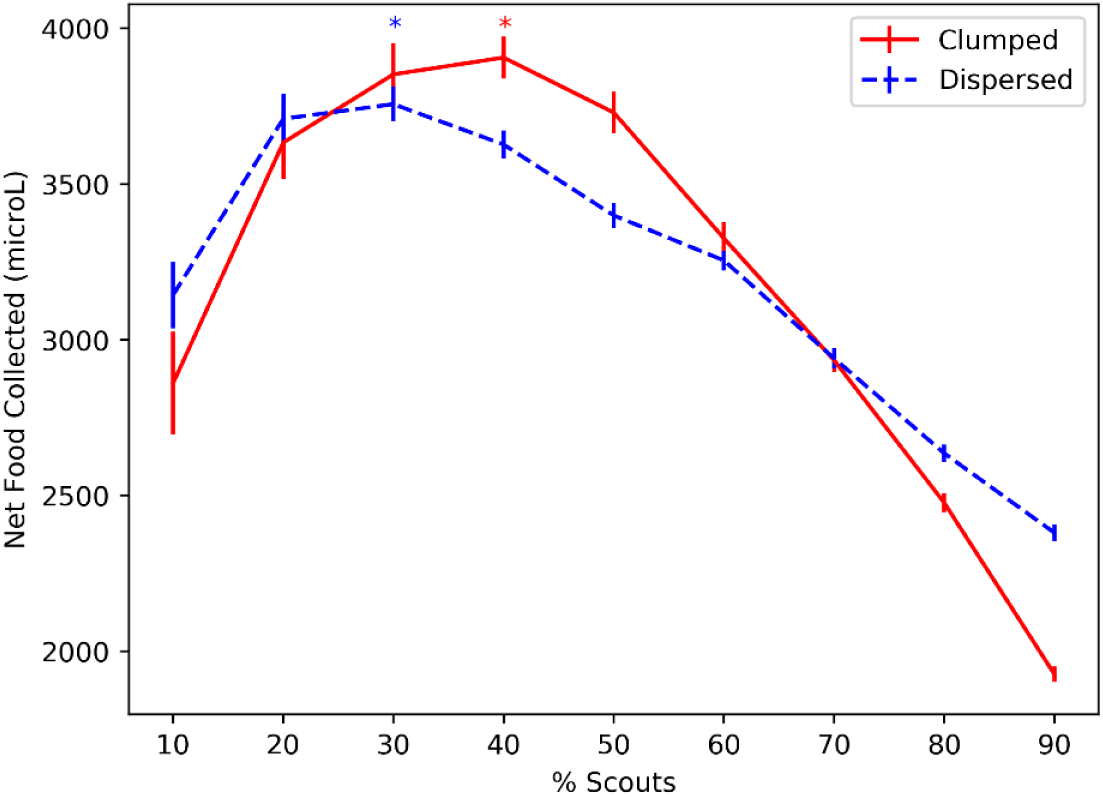
Effect of resource distribution on resource collection by simulated colonies with different proportions of scouts. Net food collected was defined as amount of food collected at the end of the simulation minus energy expended from foraging on clumped resources (red solid) or dispersed resources (blue dashed). Asterisks indicate the optimal proportion of scouts for each distribution. Vertical lines are standard errors of 150 simulation runs.

### Empirical experiments

High and low LI colonies differed significantly in their response to feeder quality (Figure 5). Low LI colonies strongly preferred to visit and consume more food from higher quality feeders over lower quality feeders, regardless of feeder distribution (GLMM: colony LI x quality; visitation: Χ^2^ = 292.01, p<0.001; consumption: Χ^2^ = 166654.01, p<0.001). In contrast, high LI colonies showed a weak preference for visiting higher quality feeders when the feeders were clumped but not when the feeders were dispersed and they did not differ in their food consumption from different quality feeders in either resource distribution (GLMM: colony LI x feeder distribution x quality; visitation: Χ^2^ = 17.37, p <0.001; consumption: Χ^2^ = 4928.69, p<0.001). Colonies from low LI lines also showed higher overall foraging activity than colonies from high LI lines, as evidenced by higher visitation and food consumption rates (GLMM: colony LI; visitation: Χ^2^= 27.10, p<0.001; consumption: Χ^2^ = 180542.3, p <0.001).

**Figure 5.**
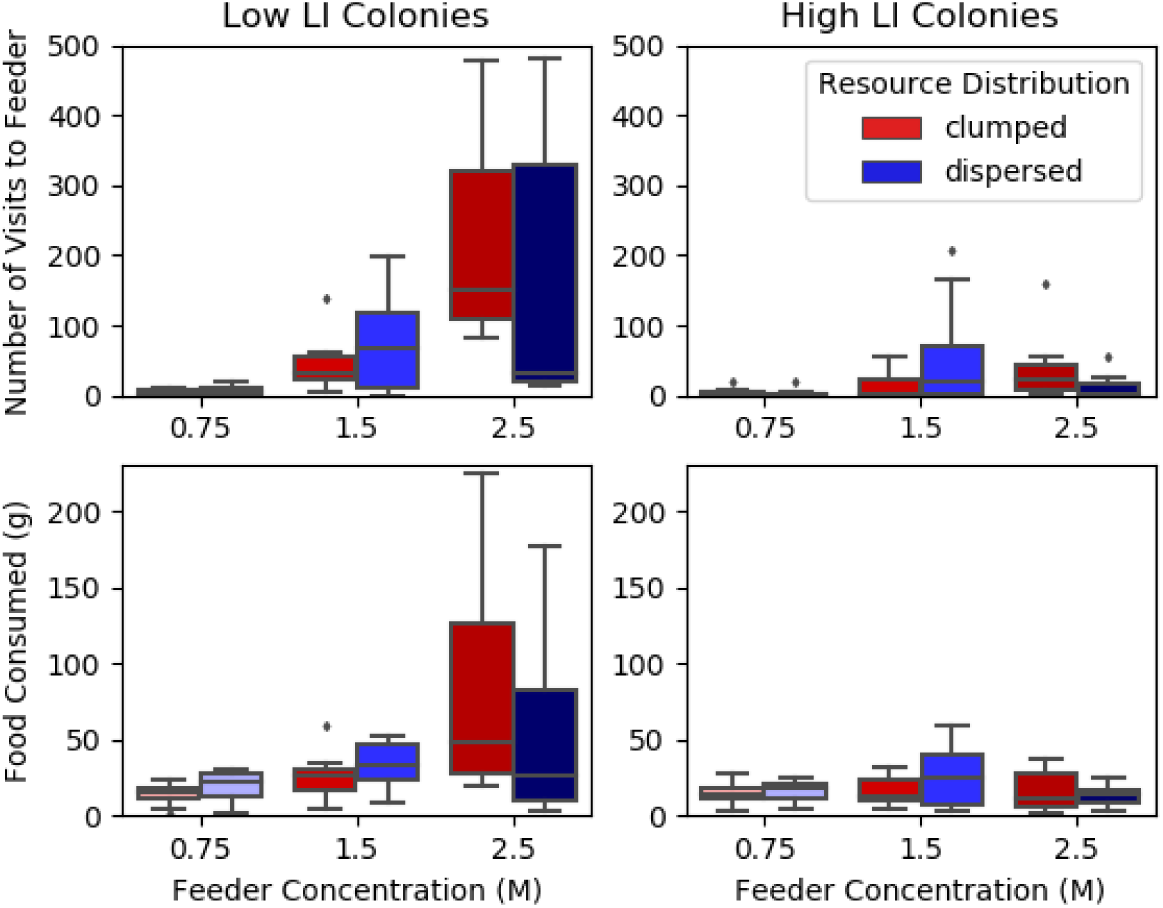
Foraging behavior of experimental colonies. Number of visits by foragers (top) and amount of food consumed (bottom) per day by colonies artificially selected for low (left) or high (right) latent inhibition (LI). Resources were clumped (red) or dispersed (blue) and varied in quality, with darker colors indicating higher sucrose concentration.

Furthermore, high and low LI colonies differed significantly in how resource distribution affected foraging behavior (Figure 5). Low LI colonies visited feeders more frequently and consumed more food when feeders were clumped compared to when they were dispersed. In contrast high LI colonies visited feeders more frequently and consumed more food when feeders were dispersed compared to when they were clumped (GLMM: colony LI x feeder distribution; visitation: Χ^2^ = 4.17, p = 0.041; consumption: Χ^2^ = 7511.26, p<0.001). For full model output see Supplementary Tables S1 and S2.

The weight lost due to evaporation in control feeders was very small relative to the amount removed by honey bees in the experimental feeders. Still, higher sugar concentration had a significantly slower rate of evaporation in the control feeders (concentration: Χ^2^ = 4.62, p = 0.03), as would be predicted by Raoult’s law [27]. Because the relationship between evaporation and sugar concentration was opposite to the relationship between concentration and food consumption observed in our experimental feeders, evaporation most likely caused us to underestimate differences in forager consumption between the different concentrations.

## Discussion

Contrary to our initial intuition, that colonies with greater investment in exploration would be better at choosing the highest quality resource patch, our simulations and empirical work show that investment in exploitation, rather than exploration, improved utilization of high quality resources. Our model predicted that colonies with greater investment in exploitation would be better at choosing the best quality resource patches and our experimental results confirmed this prediction. Colonies from the low LI line were better than colonies from the high LI line at focusing their foraging effort on higher quality patches (Figure 2, 5).

Our finding that higher investment in exploitation allows colonies to select higher quality resources suggests that collective outcomes are not a simple outcome of the composition of a group. When resources are variable in quality, organisms must not only locate resource patches, but also decide when to accept patches they find and when to continue searching for better patches. We therefore expected, initially, that high LI would help scouts to choose the best resource for their colony to exploit, because they are more selective in their learning as individuals. However, we instead found that more selective (high LI) individuals resulted in a group that is, paradoxically, less selective at the collective level.

This apparent contradiction may be explained by the nature of collective decision making in honey bee colonies. Each individual bee usually inspects only one available option and recruits other individuals to that option in proportion to its quality [15,17,28]. The colony’s choice of which resource to exploit emerges from the differences in recruitment among patches, rather than any individual directly comparing options [29–31]. High LI individuals are more likely than low LI individuals to scout independently [23], instead of relying on social recruitment to find patches. As long as a colony had sufficient scouts to locate high-quality patches, greater reliance on social information by exploiters resulted in greater utilization of the most rewarding patch by the colony as a whole and less recruitment to the least rewarding patch. *Thus paradoxically, high LI individuals, despite being “choosier” at the individual level, result in a less “choosy” collective*.

Our results highlight a key difference between the exploration-exploitation trade-off in social animals and in solitary animals. While both face a trade-off between searching for new resource patches and exploiting known patches, in social groups, individuals may choose between exploring the environment for new patches or choosing among the patches advertised by nestmates. While we initially thought of high and low LI bees as explorers and exploiters, it may be more accurate to describe them as *finders* and *refiners*: the high LI finders exploring outside the nest for available resources to advertise and the low LI refiners exploring within the nest for the best recruitment dance to follow. A colony needs enough searchers to locate available options, but the refiners are the ones who choose among them, so the colony can exploit the highest quality patch.

Our simulation model predicted that the optimal investment in exploration should be higher when resources are clumped than when they are dispersed (Figure 4). However, in contrast to this prediction, our empirical work showed that low LI colonies performed better when resources were spatially clumped, while high LI colonies performed better when resources were evenly dispersed resources (Figure 5). Characterizing high and low LI individuals as finders and refiners, instead of explorers and exploiters, may also explain why the colonies of low LI bees collected more food in the clumped resource distribution while colonies of high LI bees collected more in the dispersed distribution. A broad search pattern such as a Levy-flight is most effective for finding randomly dispersed resources [32,33], whereas when resources are clumped, individuals that are recruited to one known patch are able to also detect nearby patches.

Previous work has shown that collective investment in scouting interacts with individual persistence to influence foraging performance [8]. Because high LI individuals are more likely to scout than be recruited [23], they contribute to exploration by searching independently for food, but they can also contribute to exploitation by persisting to the sites they locate, similar to a solitary forager that learns about the environment from its experience. In contrast, low LI individuals are more likely to rely on social recruitment to find and exploit known patches. The finding that colonies with low LI bees performed better with clumped resources is therefore consistent with previous theoretical and empirical work suggesting that the benefit of recruitment information in honey bees and other social insects is greatest when resources are clumped in large patches rather than evenly dispersed [9–12].

### Conclusions

Because resources are often patchily distributed and variable in quality, gathering information about the environment is a key component to foraging success. Our results suggest a novel solution to the exploration exploitation trade-off in social groups through a division of labor between finders, who explore broadly for available resources, and refiners, who choose among discovered resources to allow the group to focus its exploitation on the most rewarding patches. Finders and refiners may differ in how they learn about resources. However the collective response to the environment is not simply an additive property of individual learning, but an emergent property of the interactions among these individuals.

## Methods

### Agent-Based Model

To examine how the distribution of food resources in the landscape influences the exploration-exploitation trade-off, we developed a spatially explicit agent-based simulation of colony foraging behavior as a modification of the model by Mosqueiro et al. [8].

To examine how colonies choose among different quality resources, we simulated a resource landscape, represented as a 36 × 36 m, 2-dimensional grid, with the hive at the center. The landscape had three 5.76 × 5.76 m resource patches, each located 14.4 m from the hive. Each resource patch had a different quality *q*, defined as the amount of food bees collected in a single foraging load (Figure 1A). The low-quality patch offered 1 resource unit per load, the medium-quality patch offered 2 units, and the high-quality patch offered 3 units. For bees, this is analogous to nectars with different sugar concentrations because load size is limited by the volume of a forager’s crop, where it carries the nectar load [34].

To uncover the effect of resource distribution on colony foraging, we simulated two different spatial distributions. In the “dispersed” distribution, the three resource patches were evenly spaced in a circle around the hive, 120° from each other. In the “clumped” distribution, all three resource patches were adjacent but not overlapping, with the centers of the patches 24° from each other (Figure 1A).

Simulated colonies contained two types of foragers: scouts, which search for food independently, and recruits, which wait to be recruited to food sources by nestmates. Forager flight dynamics were modeled as a random walk with drift [8,33]. At the beginning of the simulation (t=0), all foragers started at the hive. At t=1, scouts left the hive in a random direction and continued flying until they found food or reached the end of the foraging arena, at which point they returned directly to the hive, as in [8].

Upon returning to the hive, scouts recruited inactive foragers (“recruits”) with rate *q*γ, where *q* is the quality of the located resource and γ is a baseline recruitment rate. Scouts remained at the hive recruiting for 50 time steps and then either returned to the located resource or left the hive in a new random direction. Recruits remained inactive in the hive until recruited to a resource by another forager. Once recruited, recruits left the hive in the advertised direction and flew until they found food, at which point they returned to the hive. On returning to the hive, recruits also recruited inactive foragers at the same rate as returning scouts, *q*γ, and then either returned to the located resource or became inactive at the hive until recruited again. Both scouts and recruits returned to a known resource until their number of trips to that location was equal to their persistence parameter; see [8] for an exploration of how variation in persistence affected simulation results. Each simulation ran for 21000 time steps, equivalent to 7 hours of simulated time.

To examine how the proportion of scouts in the colony and resource distribution jointly affected collective foraging, we varied the ratio of scouts from 10%-90% of the foragers in the colony. For each scout proportion we simulated both dispersed and clumped resource distributions. Colonies were always composed of 100 foragers; see [8] for a discussion of the effects of different colony sizes. For each ratio-distribution combination, we performed 150 simulation runs. To assess how the proportion of scouts and resource distribution jointly influence collective foraging, for each run, we calculated the number of visits foragers made to each feeder as well as total food collected by the colony at the end of the simulation. We also calculated net food collected, defined as total food collected minus energy expended by foragers; see [8] for details. Finally, for each resource distribution, we calculated the optimal proportion of scouts as that which resulted in the highest net food collection.

### Empirical experiments

#### Genetic Line Selection

To create genetic lines selected for high or low latent inhibition, we reared queens by grafting 1-day old larvae into queen cups and placing them into a queenless colony with nurse bees (“queenbank”). After emergence, we placed queens into cages and back into the queenbank for 7-10 days to mature. To obtain drones for the line selection procedure, we collected mature drones as they returned to a colony from mating flights in the late afternoon and isolated them in mesh cages inside the queenbank overnight. We tested both queens and drones for latent inhibition using the procedure described below, individually marked them, and returned them to the queenbank to await insemination for no longer than 2 days.

#### Latent inhibition procedure

We scored the latent inhibition of queens and drones using a proboscis extension reflex (PER) conditioning protocol [35]. We secured individuals in a plastic harness, so that they could only move their antennae and proboscis. To ensure that these bees respond to sucrose, which is essential for the PER protocol, we presented each bee with a drop of 1M sucrose to the antennae and discarded any individual that did not extend its proboscis. We then fed each bee 7 µL of 1M sucrose and allowed it to acclimate to the apparatus for 30 minutes. We familiarized bees to one of two odors (2-octanone or 1-hexanol), both readily learned by honey bees [36], by presenting each bee with forty unreinforced 4-second bursts of odor at five-minute inter-trial intervals. To test the effect of familiarization on subsequent reinforced learning, we allowed bees to rest for 30 minutes, and then exposed them to either the familiar odor or to a novel odor, four times each, in a pseudorandom order. Both odors were equally reinforced as follows. We presented each odor for 4 seconds and if the bee extended its proboscis in the first 3 seconds, we recorded it as a positive response. Upon extending its proboscis or after 3 seconds, we rewarded the bee with a 0.4 µL droplet of 1.5M sucrose directly to its proboscis [37].

We calculated LI scores as (# positive responses to the novel odor + 1)/(# positive responses to the familiar odor + 1) and classified individuals with scores greater than 2 as high LI and individuals with scores less than 2 as low LI. We created high and low LI lineages by instrumentally inseminating high or low LI queens with like drones. We inseminated each queen with a single drone using standard instrumental insemination procedures [38], then placed it into a nucleus colony of 5000 workers for approximately one month to build up a worker population. We then placed these colonies into standard 9-frame Langstroth hives and monitored them weekly to ensure no supersedure of the inseminated queen occurred.

#### Experimental Colony Creation

To create colonies of a single behavioral type for the experiment, we placed approximately 600 newly emerged workers, marked by queen origin, of each LI type into experimental nucleus colonies. We created four experimental colonies of high LI workers and four colonies of low LI workers. To supplement the worker population of these experimental colonies, we added approximately 600 control bees from non-selected colonies. To allow the high and low LI workers to reach foraging age we waited two weeks before beginning the experiment.

#### Data Collection

To determine the foraging behavior of the selected colonies in environments with different resource distributions, we allowed colonies to forage in a controlled environment. We collected data over a two-week period, from October 1-12, 2018. We tested four colonies each week: two high LI and two low LI. Overnight, we placed each colony into the center of a 30m x 108m mesh flight tent. We allowed colonies to acclimate to the tents for one day with access to water. Due to adverse weather on the first week of the experiment, we allowed colonies tested on that week to acclimate to the tents for two days. After acclimation, to induce foraging behavior and to allow new foragers to learn how to handle artificial feeders, we provided colonies with artificial feeders containing 1 M sucrose solution scented with geraniol 2 m from the hive for one day.

To assess colonies’ abilities to choose among different quality food sources we placed three feeders of different ‘quality’ in each tent on each day of the experiment. Each feeder contained 100 mL of sucrose solution. The high-quality feeder contained 2.5M sucrose, the medium-quality feeder contained 1.5 M sucrose, and the low-quality feeder contained 0.75 M sucrose (Figure 1B). We paired each quality feeder with a unique color/odor combination. The odors used were 2-octanone, 1-hexanol, and acetophenone. Previous experience suggests no innate difference in attractiveness of the odors used [36]. Still, to control for possible differences in attractiveness, we gave each set of high and low LI colonies a different quality-color-odor combination. Each colony experienced the same quality-color-odor combination throughout the course of the experiment.

To manipulate the distribution of resources, we tested two feeder configurations, clumped and dispersed. For one high LI and one low LI colony, we placed the experimental feeders in a clumped configuration, with all three feeders closely spaced in a single corner of the flight tent (Figure 1B). For the other pair of high and low LI colonies, we placed the feeders in a dispersed configuration, with each feeder in a different corner of the flight tent (Figure 1B). To avoid biasing foragers towards a particular direction, all feeders in the dispersed configuration were in different corners from the feeders in the clumped configuration (Figure 1B). After two days, we switched the feeder distributions so that colonies that first received the clumped treatment received the distributed treatment and vice versa and collected data for two more days. To evaluate the colonies’ foraging behavior, we recorded the number of foragers that visited each feeder every ten minutes for 7 hours starting approximately at 9:00 am. We only counted foragers as visitors if they landed on the part of the feeders where food was accessible.

To measure the amount of food consumed, we weighed each feeder upon deployment and every 30 minutes throughout the experiment. We calculated daily food consumption as the difference between initial and final feeder weight each day. To determine if evaporation had a different effect on the different sucrose concentrations, we placed three feeders containing 2.5, 1.5, and 0.75 M sucrose solution each in a separate mesh enclosure from which bees were excluded. These control feeders experienced similar temperature and light conditions as the experimental ones. We quantified evaporation from each control feeder as the difference between the weight when a feeder was deployed and its weight 7 hours later.

We performed all experimental work at Arizona State University’s Honey Bee Research Lab on the Polytechnic campus in Mesa, Arizona.

#### Data Analysis

To determine how colony latent inhibition and resource distribution jointly influenced the colonies’ visitation to different quality feeders, we performed a Generalized Linear Mixed Model (GLMM) with total number of visits per day as the response variable with a Poisson log link function. We performed a second GLMM with daily food consumption as the response variable with a normal distribution and a log link function. In both models, we included colony latent inhibition, resource distribution, and feeder quality as fixed effects. We included colony ID and date as random effects to account for variation in colony strength and weather conditions respectively. We fit both models by maximum likelihood using Laplace approximation and the “bobyqa” optimizer. Visual inspection of the residuals revealed no deviation from normality. We performed all analyses in R, using the ‘lme4’ package [39]. To determine the confidence in our estimates, we performed a Type II Wald chi square test on the GLMM results, using the Anova function in the R package ‘car’ [40].

To examine whether the three concentrations differed in evaporation rates, we performed a linear mixed model (LMM) with weight lost from the control feeders as the response variable, concentration as a fixed effect, and date as a random effect.

## Acknowledgements

We thank Alexa Phillips and Eda Sezen for their help collecting and entering data, Thiago Mosqueiro for help debugging code, the Pinter-Wollman lab for suggestions throughout the preparation of the manuscript, and Matina Donaldson-Matasci for advice on methods. Funding for this work was provided by NIH grant R01GM113967.

## Author contributions

NL conducted the theoretical work, analyzed the data, and wrote the first draft of the manuscript. NL and NPW designed the simulation and outlined the manuscript. NL, NPW, CNC, and BHS designed the empirical experiments. NL, and CNC carried out the empirical experiments. CO established and maintained the bee lines. All authors commented on the manuscript.

